# Ultrasensitive quantitative protein detection using Eu-ion doped vanadate nanoparticles

**DOI:** 10.1101/2025.06.03.657629

**Authors:** Robin Kuhner, Christophe Cardone, Rafael Vieira Perrella, Fanny Mousseau, Rabei Mohammedi, Jean-Marc Sintes, Christine Bourgeois, Olivier Lambotte, Thierry Gacoin, Cedric I. Bouzigues, Antigoni Alexandrou

## Abstract

Disease prevention, diagnosis, and treatment monitoring often require ultrasensitive (sub-)femtomolar biomarker detection and quantification. While standard ELISA assays yield picomolar sensitivity, existing ultrasensitive approaches reach fM, aM or even zM sensitivity. This, however, is obtained at the expense of increased complexity and cost which hampers their biomedical applications. We propose a novel approach, NLISA, combining ultrasensitive, fM/sub-fM, quantitative detection with simplicity and ease of use based on 38-nm YVO_4_:Eu (20%) crystalline nanoparticles used as detection probes. These particles possess an extremely strong absorption in the UV leading to bright Eu^3+^-ion emission. We developed a transportable, multi-well plate reader providing LED excitation and detection with a photomultiplier enabling detection down to 16,000 nanoparticle probes/well. We obtained sensitivity gain factors with respect to ELISA ranging from 65 to 35,000 for insulin, IFN-γ, and HIV-GAG-p24 while maintaining the same antibodies. We demonstrated femtomolar LOD and a dynamic range of 4-5 orders of magnitude and NLISA efficiency for HIV-positive patient diagnosis. This approach for straightforward, ultrasensitive polypeptide/protein detection is easily generalizable paving the way for a new generation of diagnostic tests.

---

Efficient disease prevention, health care, and personalized medicine require increasingly sensitive detection systems ^1^. Indeed, the detection of lower biomarker concentrations may allow earlier disease detection and thus implementation of treatment before irreversible impact sets in and subsequently improvement of the prognosis for the patients. Ultrasensitive detection may also imply performing diagnosis without need for invasive biopsies. In particular, in the field of neurodegenerative diseases, increase of sensitivity has led to the detection of biomarkers in blood, where biomarker concentration is lower, rather than in cerebrospinal fluid requiring invasive biopsy.^2^ Moreover, ultrasensitive detection for proteic biomarkers in the (sub-)fM range promotes the identification and/or democratization of new biomarkers for more general clinical use.

Currently, in the field of protein/peptide detection, the gold standard technique is often still the enzyme-linked immunosorbent assay, ELISA, with pM detection capability: on multiwell microplates, antibodies are fixed at the bottom of each well to recognize a target biomolecule in a complex sample. The presence of the target or analyte is then revealed by a second antibody functionalized with a label, an enzyme producing an absorbing chromophore in the presence of a substrate. ELISA variants like electrochemoluminescence immunoassays (ECLIA) yield increased sensitivity. ECLIA uses a revelation system based on the luminescence properties of a ruthenium complex which is excited by a current *via* excitation transfer from a sacrificial partner.^3,4^ Indeed, detection approaches using light emission are generally more sensitive than those based on absorption because the signal detection takes place in background-free conditions whereas absorption signals are inherently detected in the presence of the incident light background. The obtained typical 0.1 pM protein detection capability is, however, often not sensitive enough.

The ELISA detection sensitivity could be enhanced through amplification approaches. Using DNA labels instead of enzymes and combining ELISA with qPCR yields quantitative sensitive immune-PCR^5,6^. Alternatively, a barcode approach combining oligonucleotide labelling, their amplified release and detection on DNA chips was proposed by Mirkin and coworkers^7^. Both approaches exploit the efficacy of nucleic acid detection techniques and reach extreme aM sensitivities but at the expense of complexification and lengthening of the detection process.

More recently, single-molecule counting based ELISA has reached exquisite sensitivity either by capturing and detecting enzyme-labeled antibodies in patterned substrates with multiple wells through a sensitive 2D camera^8–10^ or by detecting fluorescence bursts of single analytes immuno-labeled by fluorophores on a confocal set-up.^11^ An aM-sensitivity has been demonstrated for very long incubation times^8^ and may be extended to the zM range by dramatically increasing the number of wells in the case of a patterned substrate.^9^ However, in both cases, complexity and cost limit their clinical application for disease diagnosis or monitoring.^12^ Indeed, commercialized realizations of these technologies by Quanterix and Singulex, respectively, enable fM sensitivity but remain limited to research use.

Other ultrasensitive polypeptide detection approaches, designated plasmonic ELISA, exploit the shift of the surface plasmon resonance to reach sensitivities in the fM, aM or zM range. Notably, the inverse sensitivity approach uses the enzyme glucose oxidase producing H_2_O_2_, which in turn controls the deposition of a Ag layer on Au nanostars, as the detection label^13^. Alternatively, the enzyme catalase degrading H_2_O_2_, which controls the formation of Au nanoparticles, is used as the labelling molecule.^14^ These techniques are complex and rely on multiple components: enzymes whose activity may show batch-to-batch variability and require freezing, metallic ion solutions which need to be freshly prepared, and nanoparticle synthesis reactions sensitive to parameters like pH and temperature. Most importantly, these approaches are not quantitative.

Plasmonic metal nanostructures have been used for localized enhancement of the electromagnetic field under laser excitation to yield Surface-enhanced Raman spectroscopy (SERS)^15^ enabling detection of the specific Raman signal of biomolecules^16^ or pathogenic organisms like bacteria^17^ down to the single biomolecule/organism. Nevertheless, SERS requires complex instrumentation. Further approaches include temperature-responsive dye-containing liposomes as probes reaching 970 zM sensitivity, however without quantitative detection ^18^.

Overall, current approaches to extend sensitivity to the (sub-)fM range suffer from common limitations: the gain in sensitivity comes at the expense of complexity and cost increase, which often hampers their transfer to commercial and clinical use. Moreover, although experimental demonstrations of aM-zM sensitivity represent exciting scientific achievements, the main need in *in vitro* diagnostics concerns the 0.1 fM-1 pM range.

In this context, we propose the use of YVO_4_:Eu nanoparticles as luminescent revelation probes in an ELISA-like scheme, combined with an *ad hoc* optical detection system. This approach benefits from the unique optical and colloidal properties of these lanthanide-ion doped vanadate nanoparticles. YVO_4_:Eu nanoparticles, when excited through the vanadate matrix, present an extraordinary brightness resulting from the strong vanadate absorption coefficient and a good quantum yield. They yield large luminescence signals detectable with a simple photomultiplier (PMT) even when present in small numbers thus pushing ELISA-like protein detection from the pM range to the (sub-)fM range of ultrasensitive single-molecule-based detection without any increase in instrumentation complexity or cost. We have implemented it in a 96-well plate, transportable reader using simple, inexpensive UV LED excitation, compatible with use in high- or low-technology environments. We designate this approach NLISA for Nanoparticle-Linked ImmunoSorbent Assay.

## Results

### Luminescent YVO_4_:Eu nanoparticle synthesis and functionalization

Lanthanide ions, in particular Eu^3+^ ions, are well known for their luminescent properties and have been synthesized in particle form in a variety of host compounds^19^. Key optical properties of such Eu-doped nanoparticles include: i) narrow emission, ii) large Stokes shift with respect to the excitation wavelength allowing efficient rejection of background light, iii) high photostability and absence of blinking, and iv) long luminescence lifetime (a few hundreds of µs) which can be exploited for time-resolved detection^20^. Direct Eu^3+^-ion excitation is possible at 466 or at 396 nm and 30-nm nanoparticles containing several tens of thousands of ions exhibit absorption and luminescence signals similar to those of good organic fluorophores.^19^ A matrix consisting of vanadate ions offers the additional advantage of introducing a strong charge-transfer absorption peak at 280 nm compatible with subsequent energy transfer to the europium ions increasing the absorption coefficient by more than three orders of magnitude. Note that the nanoparticle brightness is determined by its size and cristallinity, because both the absorbing and emitting ion numbers scale with the particle volume.

Colloidal YVO_4_:Eu 30-40-nm nanoparticles were synthesized by a room-temperature aqueous co-precipitation approach developed by Huignard *et al*^21,22^. This yielded nanoparticles with sizes and brightness compatible with single-particle detection^19,23,24^ and sensing^25–27^ under direct excitation of the Eu^3+^ ions with an epifluorescence microscope. UV excitation (∼280 nm) of the vanadate matrix is harmful for living cells and is expected to induce strong autofluorescence signals, even for *in vitro* applications, hindering high sensitivity detection. First efforts to obtain ultrasensitive detection were thus focused on direct Eu^3+^ excitation using a 1-W 466-nm diode laser^28^. We nevertheless show below that the strong autofluorescence background due to UV excitation is overcome by the efficacy of the nanoparticle absorption at the vanadate absorption peak with the additional advantage that a 100-mW inexpensive LED could be used.

Here, we used an improved YVO_4_:Eu nanoparticle synthesis exploiting a metavanadate ion (VO_3_^-^) instead of an orthovanadate ion (VO_4_^3-^) precursor to obtain particles of controlled size and crystallinity^29^. The crystallinity was then further improved (Fig. 1A-B) to increase the quantum yield by a post-synthesis heating protocol at 200°C (Materials and Methods). In these crystalline particles, improved energy transfer processes in the vanadate sub-lattice allow optimized emission at lower Eu content, thus limiting activator concentration quenching phenomena.^30,31^ Therefore, a doping level of 20% is used, lower than the 40% doping in non-annealed less crystalline nanoparticles previously used^30,31^. The quantum yield of the obtained YVO_4_:Eu (20%) nanoparticles was 18%. The particles were then functionalized using previously published protocols involving surface silicatation and silanization^32–34^ and coupled to streptavidin molecules, before conjugation to biotinylated antibodies (Materials and Methods). Even though it is possible to directly conjugate functionalized nanoparticles to the antibodies of interest, this approach is used for higher versatility.

**Fig. 1.**
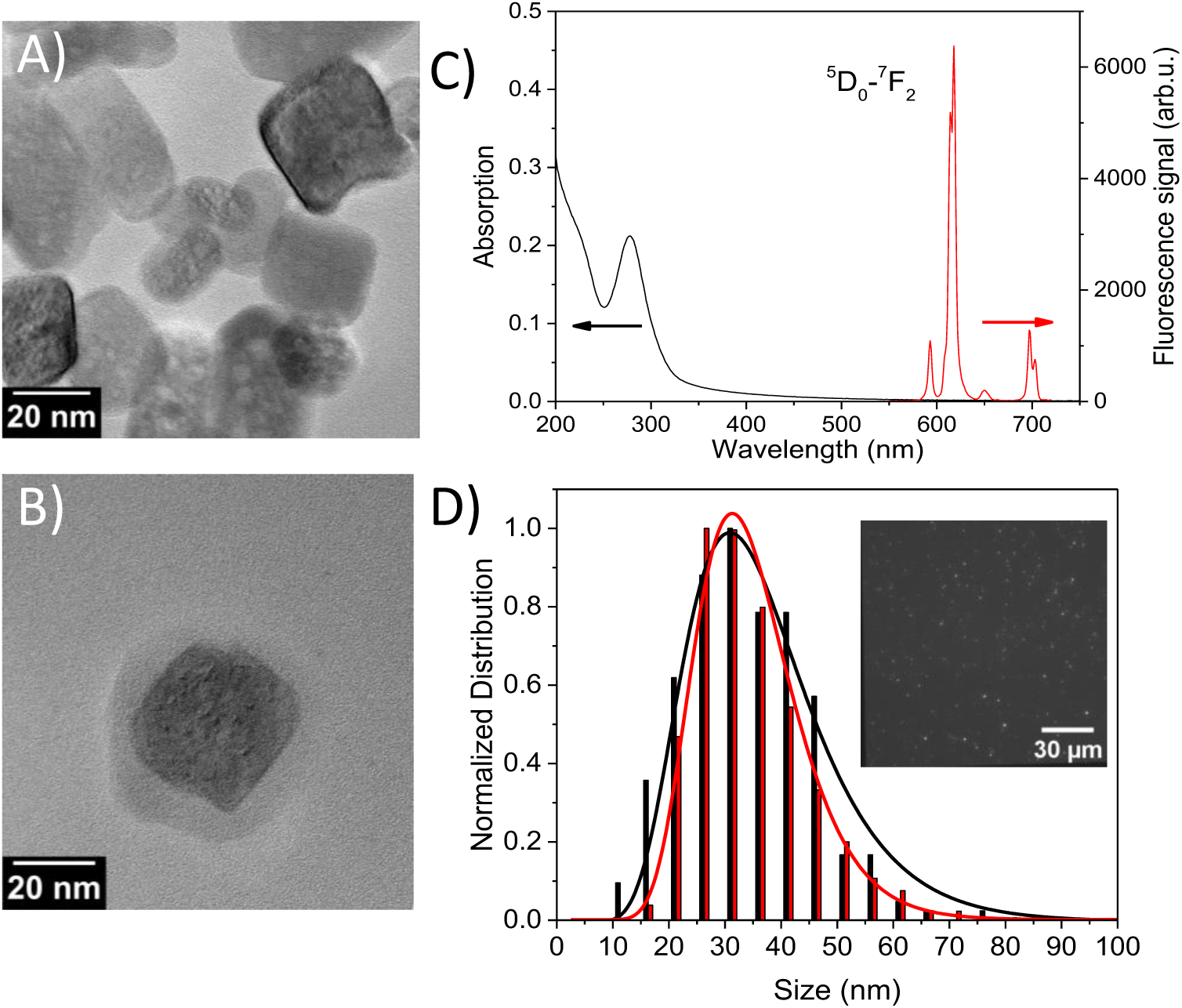
Characterization of annealed YVO_4_:Eu 20% nanoparticles. A) TEM image of bare YVO_4_:Eu 20% nanoparticles. B) TEM image of a silica-coated YVO_4_:Eu 20% nanoparticle. C) Absorption (black) and emission spectra (red; excitation wavelength: 280 nm) of silica-coated annealed YVO_4_:Eu 20% nanoparticles. D) Size distribution histograms of silica-coated YVO_4_:Eu 20% nanoparticles determined from TEM images (black; N=262) and from the number of detected photons under 466-nm direct Eu^3+^-ion excitation using a wide-field inverted microscope (red; N=3633). The solid lines are log-normal fits of the two distributions. The inset of D) shows a typical optical microscopy image exhibiting numerous single-nanoparticle emission spots.

### Nanoparticle characterization

The nanoparticles were characterized using transmission electron microscopy (TEM) before and after coating with a silica layer that stabilizes the particles and facilitates further functionalization (Fig. 1A and B, respectively). The mean nanoparticle volume and the corresponding size (diameter of a sphere with the same volume as the particles) were determined from the TEM images to be *V*_*NP*_ = 28 900 nm^3^ and *D* = 38 nm, respectively (Fig. 1D). Given that the unit cell volume of YVO_4_:Eu (20%) is *V*_𝑢𝑛𝑖𝑡_ = 0.32 nm^335^ and that the unit cell contains 4 vanadate ions, the mean number of vanadate ions per nanoparticle is 360 000. We can then estimate the remarkably high extinction coefficient of these nanoparticles at 280 nm 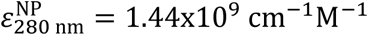 from the extinction coefficient of a vanadate ion at 280 nm 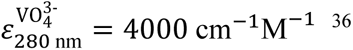 ^36^ (to be compared to nanoparticle extinction coefficient upon direct excitation of Eu^3+^ ions at 466 nm for a 20% Eu doping, 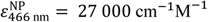. These nanoparticles thus exhibit extreme brightness of 2.6×10^8^ cm^-1^.M^-1^. Therefore, strong luminescence signals can be obtained with low excitation intensity. Absorption and emission spectra of these nanoparticle solutions are presented in Fig. 1C.

As previously demonstrated for non-annealed YVO_4_:Eu nanoparticles^20^, we confirmed that the TEM-based size distribution is preserved in optical imaging conditions and is in good agreement with the size distribution obtained by the analysis of the photon distribution number of single nanoparticles as follows: we deposited and incubated a diluted solution of silica-coated nanoparticles on different wells of a microplate and then, after rinsing, imaged the non-specifically adsorbed particles (inset Fig. 1D, Supp. Materials and Methods). Single particles are observed as diffraction-limited emission spots (absence of aggregates). Multiple images were analyzed to yield the photon number distribution which is then converted into size distribution (Fig. 1D) using (Equ. 1):

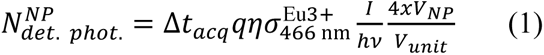

where 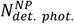 is the photon number detected/NP/s, *q* the particle quantum yield, 𝜂 = 20% the microscope detection efficiency, 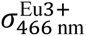 the absorption cross section of a single Eu^3+^ ion, *I*/ℎ*v* the incident photon number per unit surface, and 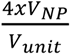 the number of Eu^3+^ ions per nanoparticle with *x* = 0.2 the fraction Eu/Y+Eu of Eu ions. The overlap between the particle size distribution obtained from TEM images and from the optical microscopy images confirms that the nanoparticles show excellent colloidal dispersion, that their dispersion state is unaffected by the deposition on substrates (for TEM or optical imaging), and that the detected photon number can be used to determine the nanoparticle size.

### NLISA implementation on a nanoparticle luminescence reader

For the readout of the NLISA assays, we developed a home-made microplate reader which provides UV excitation with a 275-nm, 100-mW LED to benefit from the strong vanadate absorption, compatible with readout of 96-well microplates (Fig. 2B, Materials and Methods, Supp. Table S1).

**Fig. 2.**
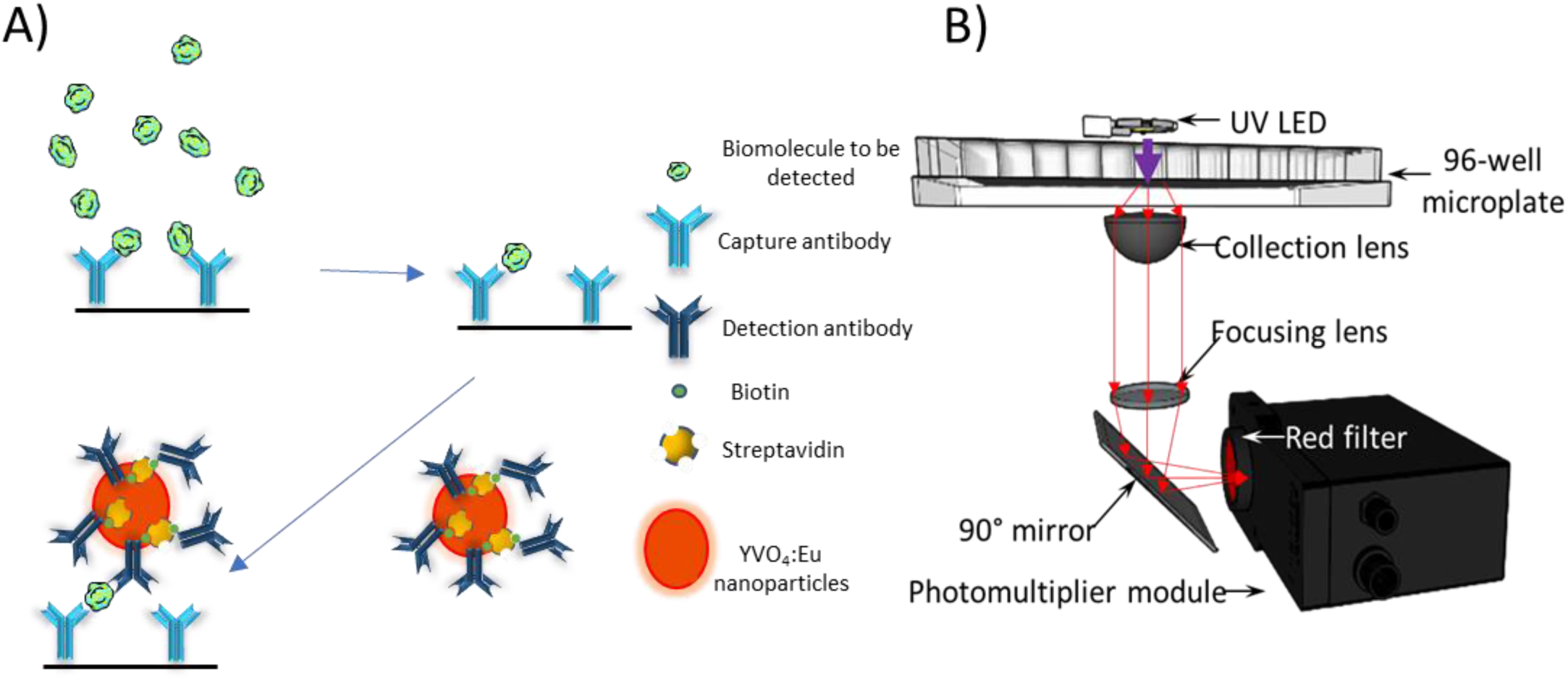
A) Schematic of the NLISA principle. A surface coated with capture antibodies is incubated with the sample containing the target protein (analyte) (top left). After rinsing, non-attached target molecules and other molecules in the sample are removed leaving the antibody-bound target molecules on the surface (top right). We then incubate with the nanoparticule-detection antibody conjugates and, after rinsing to remove unbound conjugates, the nanoparticle-labeled target molecules (bottom left) can be detected through the luminescence emission of the nanoparticles upon excitation. B) Home-made microplate reader featuring a 275-nm, 100-mW LED for the nanoparticle excitation, a diaphragm (not represented) to selectively illuminate a single well and avoid cross-talk between multiple wells (Supp. Table S1), a lens collimating the particle emission and a second lens focusing it onto a multi-alkali 300-800 nm photomultiplier tube (PMT) through two 620-nm interference filters selecting the nanoparticle luminescence.

The enzyme labels in ELISA transform a substrate into absorbing or emitting molecules. In NLISA, they are replaced by YVO_4_:Eu nanoparticles (Fig. 2A). We determined the lowest detectable analyte concentration with our setup using LOD_3σ_ (limit of detection), the concentration yielding Signal(LOD_3𝜎_) = Signal_blank_ + 3𝜎_blank_, where Signal_blank_ and 𝜎_blank_are the signal of a blank sample and its standard deviation, respectively. The Signal_blank_ results both from the background signal, due to molecules and materials in the sample or in the reader (around 100 mV), and from the non-specific adsorption of the nanoparticle probes on the detection surface (typically 50 mV for the nanoparticle concentrations used here). Thus, efficient passivation and rinsing to remove non-specifically adsorbed nanoparticles are crucial to minimize our LOD. The fluctuations of the blank signal arise mainly from well-to-well variations and thus have a biochemical origin due to the variability of non-specific nanoparticle adsorption.

We next determined the minimal number of nanoparticles necessary to obtain a detectable signal, LOD_N_, *i. e.* necessary to obtain a signal above Signal(LOD_3𝜎_). We calibrated the reader by comparing the signal due to nanoparticles obtained above the background signal in the absence of particles to the average number of single particles identified in optical microscopy images such as those in the inset of Fig. 1D (Supp. Materials and Methods) normalized to the well surface and obtained a correspondence of 1,600 nanoparticles/well per mV of reader signal. We moreover checked that the PMT response is linear (Supp. Fig. S2). For the case of IFN-γ and p24 detection (see below), 3𝜎_blank_is equal to 10 mV which yields LOD_N_ = 16,000 nanoparticles. By assuming that this nanoparticle number is equal to the lowest detectable number of analyte molecules and based on the sample volume of 100 µL used for the detection, we estimate the minimal detectable analyte concentration to be LOD_est_ ∼0.27 fM. Overall, these results demonstrate the NLISA capability of sub-fM/fM-range detection.

### Insulin detection

As a first test of the performance of our approach we chose insulin in aqueous buffer as a model analyte. To determine the gain in sensitivity with respect to the standard horseradish peroxidase-based ELISA assay, we used the same antibody pair and buffers as in a commercial ELISA test. We determined the LOD_3σ_ of the commercial test to be 87.6 pg/mL in agreement with specifications: 50 pg/mL or 4.17 pM (Fig. 3A inset). In contrast, the lowest detectable concentration with NLISA is in the fg/mL range: the *lowest detected concentration* yielding a signal above the LOD_3σ_ value is 10 fg/mL (834 aM; Fig. 3A). Moreover, we find a linear signal dependence over several concentration decades. By fitting this NLISA calibration curve with a Hill equation (n=1), we obtain the *lowest detectable concentration* as the concentration for which the Hill fit crosses the LOD_3σ_ signal value red line and find LOD_3σ_ =1.43 fg/mL (119 aM) (Fig. 3A). This means that our NLISA assay is 35,000 times more sensitive than the commercial ELISA test. This LOD_3σ_ concentration is lower than LOD_est_ and may indicate that multiple nanoparticles can bind to a single insulin molecule.

**Fig. 3.**
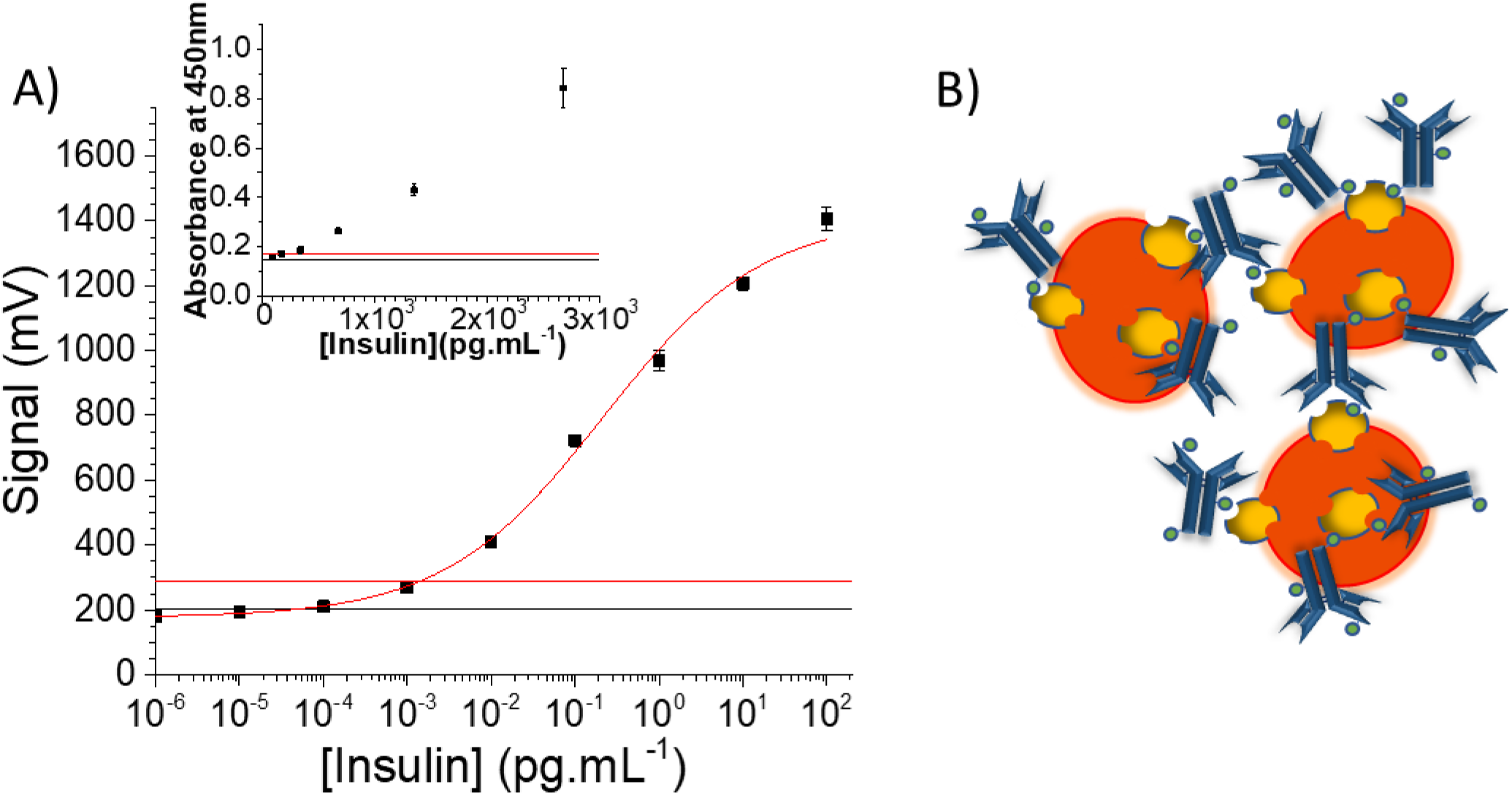
A) Insulin detection using NLISA with annealed YVO_4_:Eu 20% nanoparticles vs. the standard ELISA approach shown in the inset. The black line represents the signal value of the blank (average of 10 measurements) while the red line represents the signal value at *Signal*_*blank*_ + 3𝜎_*blank*_thus indicating the value above which a signal is considered positive with 99.7% certainty and determines the LOD_3_σ concentration. The measurements were performed in triplicate and their error bar corresponds to the standard error on the mean. The data are fitted with a Hill curve (n=1). B) Nanoparticle complex formation exploiting multiple biotins on each anti-insulin antibody proposed to explain the fact that multiple nanoparticles must be labeling each insulin molecule (Supporting Table S2).

As discussed above, we can estimate the number of nanoparticles corresponding to the various PMT signals and compare it to the number of insulin molecules in the 100 µL sample (Supp. Table S2). For concentrations at or below 8.34 fM, we obtain higher nanoparticle numbers than the maximum possible number of surface-bound insulin molecules. These results showing a nanoparticle:analyte ratio of 1.7:1 for 8.34 fM and 6.7:1 for 0.834 fM (Supp. Table S2), are indicative of the presence of nanoparticle complexes containing multiple particles (Fig. 3B) and labelling a single antigen. These complexes may originate from (i) the fact that the detection antibodies are multibiotinylated and (ii) the large number of streptavidin molecules conjugated to nanoparticles (37:1, Materials and Methods) and may significantly increase the probe signal emitted per analyte molecule. Furthermore, the formation of these complexes may enhance the effective affinity for the analyte through an avidity effect.^37–39^ In contrast, in an ELISA assay, the horseradish peroxidase-streptavidin (HRP-Strep) chimeras are intrinsically produced in a 1:1 stoichiometry, which prevents such an effect.

To further confirm the hypothesis of nanoparticle complex formation, we deposited streptavidin-functionalized nanoparticles (NP-SA) and nanoparticles conjugated to multibiotinylated insulin and IFN-γ antibodies on an olefin 96-well plate at concentrations yielding similar signal values when read with our reader. We imaged these nanoparticles under a wide-field microscope upon 466 nm excitation in the same conditions as the silica-coated nanoparticles in Fig. 2D (Fig. 4A). When imaged with a 500-ms acquisition time (Fig. 4A top row), the NP-SA conjugates (left column) are clearly observed as individual particles similarly to what is observed for silica-coated particles in Fig. 1D. In contrast, NP-SA-anti-insulin Ab and NP-SA-anti-IFN-γ Ab conjugates (Fig. 4A top center and left, respectively) appear as much brighter spots leading to saturation of the camera. To avoid saturation, the same wells were also imaged with a much lower acquisition time of 100 ms (Fig. 4A bottom row).

**Fig. 4.**
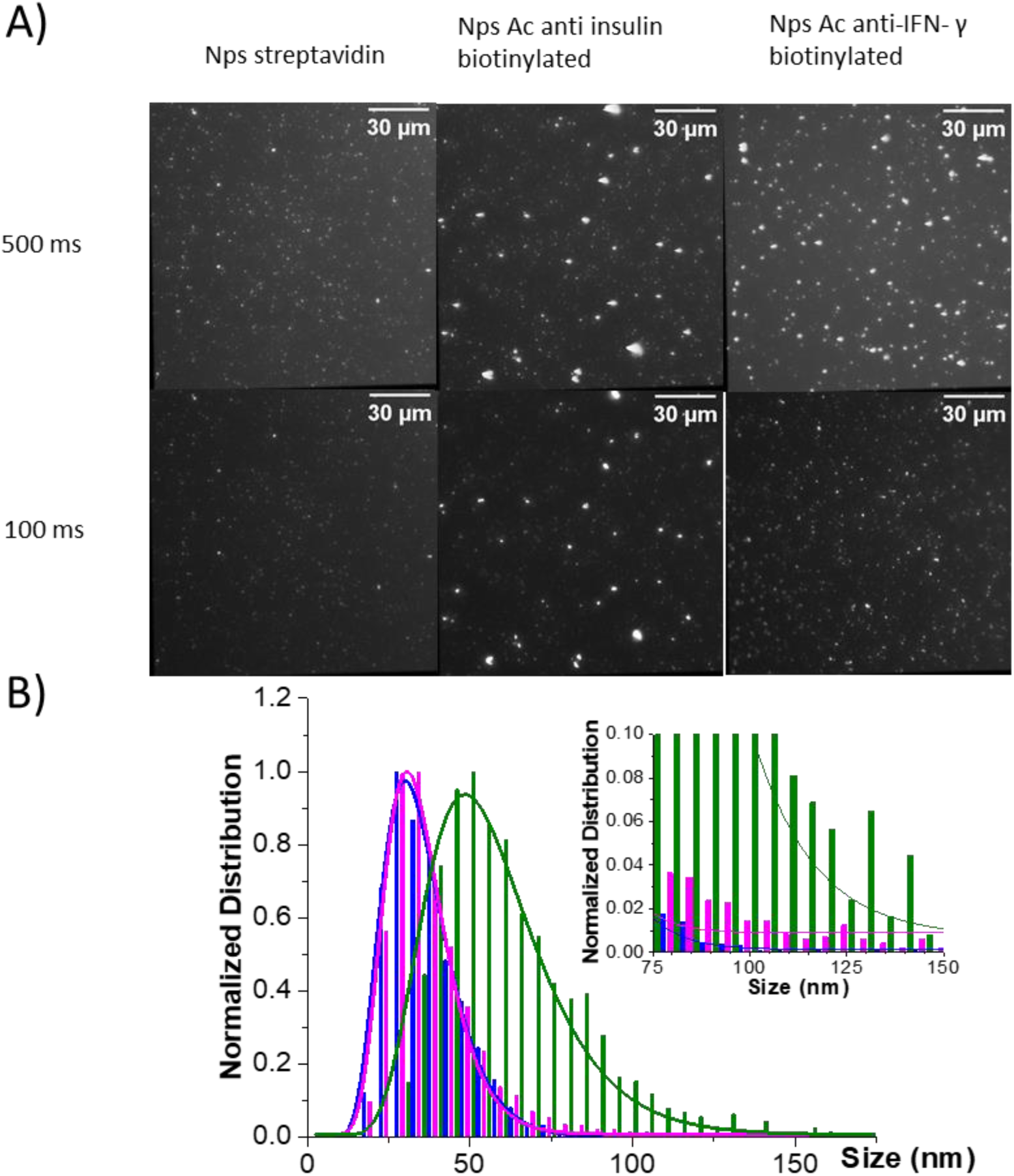
A) Wide-field microscopy images of functionalized annealed YVO_4_:Eu 20% nanoparticles in the presence or absence of biotinylated antibodies at two different acquisition times: 500 ms (top) and 100 ms (bottom) to reveal the presence of nanoparticle complexes. Left column: NP-SA conjugates. Middle: NP-SA-anti-insulin Ab conjugates. Right: NP-SA-anti-IFN-γ Ab conjugates. The concentrations were adjusted so that similar signal values are obtained on the reader PMT: 2234, 1976, and 2146 mV, respectively. The photon numbers per emission spot obtained for NP-SA-anti-insulin Ab and NP-SA-anti-IFN-g Ab conjugates for an acquisition time of 100 ms were multiplied by 5 to be directly comparable to those for NP-SA conjugates and those shown in Fig. 1D obtained for an acquisition time of 500 ms. B) Size distribution and log-normal fit of annealed nanoparticles calculated from the detected photon numbers in wide-field microscopy images (see text) for nanoparticles functionalized with streptavidin (blue N=8869), with multibiotinylated antibodies targeting insulin (pink; N=4870), and with multibiotinylated antibodies targeting IFN-γ (green; N=2092).

By analyzing multiple images acquired during 500 ms for NP-SA conjugates and during 100 ms for NP-SA-anti-insulin Ab and NP-SA-anti-IFN-γ Ab conjugates, we plotted the photon number distribution for each population. Applying equation (1), we extract the corresponding size distributions (Fig. 4B). The size distribution of the NP-SA conjugates is similar to the one for silica-coated nanoparticles (Fig. 1D) indicating the absence of size modification and aggregation effects during the functionalization steps (the two size distributions are compared in Supp. Fig. S3). In contrast, the size distribution for NP-SA-anti-IFN-γ Ab conjugates is strongly shifted to larger sizes and the distribution for NP-SA-anti-insulin Ab conjugates shows a stronger contribution of large-sized nanoparticles (inset of Fig. 4B), confirming the presence of nanoparticle complexes containing several nanoparticles each. These results confirm the formation of nanoparticle complexes leading to further enhancement of the NLISA sensitivity in the sub-fM range beyond the minimal detectable density of single nanoparticles.

### Inflammation and infection marker detection

After this proof-of-concept detection of insulin, we applied our NLISA scheme to two proteins of interest, the interferon-γ (IFN-γ), a biomarker signalling inflammatory processes which may be found in low concentrations in patient serum, and HIV-GAG-p24, the internal capsid protein of the HIV virus, to illustrate its relevance for inflammatory and infectious diseases.

IFN-γ is a cytokine strongly involved in inflammation^40^. Produced in T lymphocytes ^41,42^, it is involved in many cellular mechanisms^43–45^ such as innate or adaptive immunity^43,45,46^ and has pro- or anti-tumor effects.^47,48^ It is already used in combination with other interleukins^40^ to identify latent tuberculosis^49,50^ or other diseases but more sensitive detection would allow early management of a variety of pathologies including sepsis^51–54^.

The p24 protein is a usual target in the context of HIV diagnosis^55,56^. It is crucial for early HIV diagnosis in the context of acute infection, *i. e.* when an antibody response has not been mounted yet, and is typically used for diagnosis in conjunction with antibody detection. At later stages of the disease, anti-p24 antibodies form immune complexes sequestering p24. Detecting p24 in such situations requires immune complex disruption^57,58^ and is essential for early infant diagnosis for infants born to an infected mother where the mother antibodies present in the infant circulation prevent antibody-based detection.^59^ HIV RNA detection using qPCR is the preferred solution during therapy, where very low viral load levels need to be detected, to monitor patient adherence to the treatment or resistance appearance. Even though HIV diagnosis using qPCR is possible already few days after infection^57,60,61^, it is usually reserved to viral load measurements because of cost and complexity issues.

IFN-γ detection in buffer using a standard ELISA commercial test yielded LOD_3𝜎_ = 18 pg/mL (Supp. Fig. S4A), in agreement with supplier specifications 4 pg/mL. Using the same antibodies and buffers, our NLISA assay gave LOD_3𝜎_ = 40 fg/mL (2.28 fM) (Fig. 5A), *i.e.* a sensitivity gain of 100 with respect to ELISA. Moreover, the LOD_2.5𝜎_ = Signal_blank_ + 2.5𝜎_blank_ value is 32 fg/mL comparable to the value of 16 fg/mL^62^ obtained with the ultrasensitive single-molecule approach SIMOA© requiring much more complex and expensive equipment, consumables, and data analysis. For the lowest detected concentration (Supp. Table S3), the nanoparticle:analyte ratio is 0.08, assuming that all IFN-γ molecules are bound to the surface, assumption which is not plausible. We consider that this nanoparticle:analyte ratio may be sufficiently close to 1 to indicate the presence of nanoparticle complexes containing multiple particles (Fig. 3B and 4B).

**Fig. 5.**
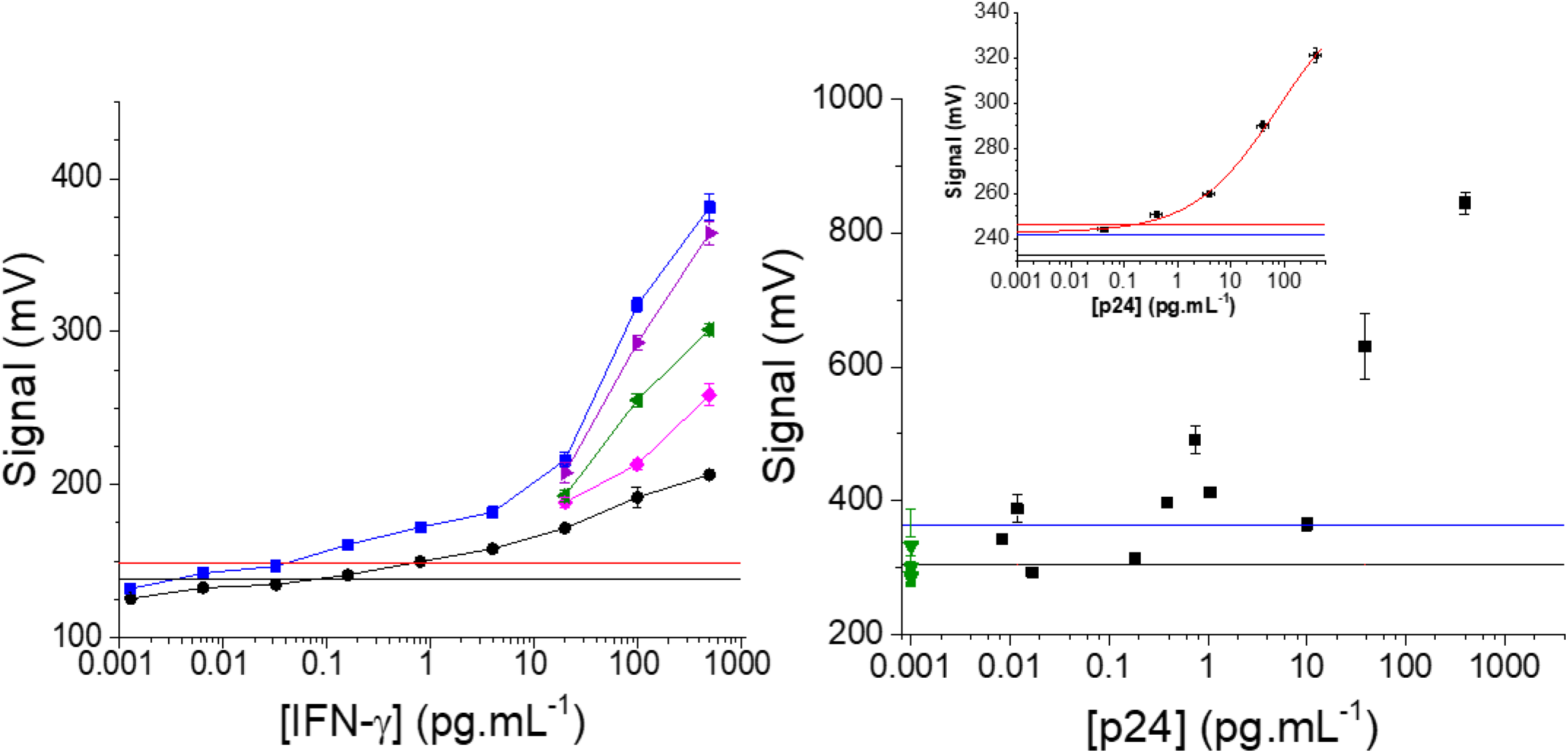
NLISA with annealed YVO_4_:Eu 20% nanoparticles. A) IFN-γ detection. Data points correspond to results in buffer (blue), in different serum dilutions, 10x (purple), 5x (green), 2x (pink), and in undiluted serum (black). The black and red straight lines represent the mean of the blank signal *Signal*_*blank*_and the signal value at *Signal*_*blank*_ + 3𝜎_*blank*_, respectively, obtained from 10 measurements. The LOD_3_σ signals (red line) in buffer or serum or diluted serum are quasi-identical and thus undistinguishable on the figure. We find an LOD_3_σ in buffer (serum) of 40 (625) fg/mL, respectively. Experiments performed in triplicate and error bars correspond to the standard error on the mean. The solid lines connecting the data points are a guide to the eye. B) Detection of p24 in patient plasma samples. In green, samples from healthy patients (n=6); in black, samples from infected patients (n=10). Experiments were performed in triplicate; error bars correspond to the standard error on the mean. The black and blue straight lines represent the mean of the negative samples and the signal value at *Signal*_*blank*_ + 2𝜎_*blank*_, 363.9 mV, respectively. The inset shows p24 detection in spiked buffer. The black, blue, and red straight lines represent the mean of the blank samples, the signal value at *Signal*_*blank*_ + 2𝜎_*blank*_, 241.8 mV, and the signal value at *Signal*_*blank*_ + 3𝜎_*blank*_, 246.4 mV, respectively, obtained from 10 measurements. The data are fitted with a Hill curve (n=1; red line). Experiments were performed in triplicate; error bars correspond to the standard error on the mean. Horizontal error bars indicate the uncertainty in the p24 concentration provided by the supplier.

When performing the same test in serum, both ELISA and NLISA sensitivity is degraded by a factor of approximately 10 (Fig. 5A and Supp. Fig. S4B). Serum constituents (proteins, peptides, …) impair antibody-antigen interactions and effectively decrease the assay sensitivity. It is therefore common for detection assays to be conducted in serum diluted with buffer. In undiluted serum spiked with IFN-γ, we find LOD_3𝜎_ = 625 fg/mL (36.5 fM), whereas successive dilutions show that detection in 10x-diluted serum is similar to the one in aqueous buffer. The ELISA assay in undiluted serum yields LOD_3𝜎_ = 42 pg/mL (Supp. Fig. S4B). NLISA thus enables a sensitivity gain of 67 in undiluted serum. Note that there is a non-zero IFN-γ concentration in sera of healthy donors of approximately 0.91±0.38 pg/mL^63^ which somewhat affects the LOD determination in serum.

A significant sensitivity gain was also observed for the p24 protein (MW=24 kD) in buffer. The LOD_2𝜎_ specification of the ELISA test is 3.25 pg/mL, whereas our NLISA approach enables a LOD_3𝜎_ between 117 fg/mL (4.8 fM) and 191 fg/mL (7.95 fM) obtained by fitting the data points with a Hill equation (n=1) with a smallest detected concentration between 307 and 500 fg/mL and a LOD_2σ_ under the last concentration measured, between 50 and 30.7 fg/mL (inset Fig. 5B). This yields a sensitivity gain between 65 and 106, compared to ELISA. The highest nanoparticle:analyte ratio calculated in the Supp. Table S4 is 0.03.

### Application to patient samples

Inactivated sera from healthy (N=5) and HIV-positive patients (N=10) were measured after 5x dilution (Fig. 5B). The negative samples show significant variability leading to an LOD_2𝜎_ signal value higher than in buffer (363 mV vs. 246 mV, inset of Fig. 5B). The concentration of p24 was independently determined by qPCR by considering that each virion contains 2,000 p24 molecules.^64^ Although four of the positive samples seem to follow a Hill-like curve, the remaining six samples yield a signal below the expected one. This is probably due to the fact that patient-secreted anti-p24 antibodies can bind p24 rendering it undetectable.^58^ Even in the presence of this variability, p24 was unambiguously detected for concentrations as low as 387 fg/mL (16.1 fM) corresponding to 4860 viruses/mL. The detection threshold is significantly lower than for spiked samples in buffer. This may originate from degradation of the HIV RNA leading to underestimate the p24 concentration through qPCR. Alternatively, multiple p24 molecules in the viral capsid may induce avidity effects absent in calibration measurements with isolated, independent p24 molecules.

## Discussion

We obtained a significant gain in sensitivity using bright luminescent YVO_4_:Eu nanoparticles as probes with respect to ELISA tests using horseradish peroxidase for three different biomolecules: insulin in buffer, IFN-γ both in buffer and serum, and HIV-GAG-p24 in buffer and in patient samples (Table 1). This sensitivity gain originates exclusively from the probe efficiency and its detection method, as we used the same antibodies and solvents as for ELISA. The sensitivity gain factor we obtain depends on the biomolecule and its specific antibodies but in all cases an LOD_3𝜎_ of a few fM or less was obtained, comparable to the sensitivity obtained by current commercial ultrasensitive technologies like SIMOA©. In the case of insulin and IFN-γ, the determination of the nanoparticle number from the reader signal lead us to propose the formation of nanoparticle complexes which further enhance the sensitivity, down to the aM range, in the case of insulin. In the case of p24, detection down to 387 fg.mL^-1^ / 16 fM (4860 viruses/mL) was unambiguously possible. The detection here is semi-quantitative due to partial inactivation of p24 by serum antibodies.^65^ Immune complex disruption may render quantitative p24 detection feasible down to 12 fg/mL (152 viruses/mL; see data point in Fig. 5B) approaching the qPCR sensitivity of a few tens of viruses/mL. Moreover, an essential feature is that these results were obtained with a simple, easy to implement protocol and a small transportable reader.

**Table 1.**
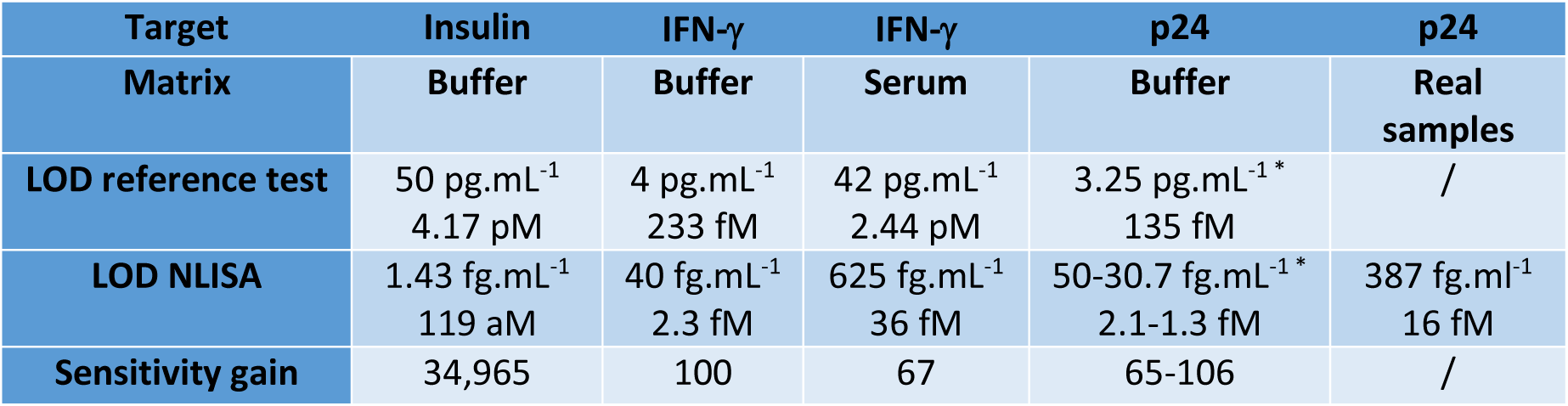
Summary of sensitivities, LOD_3_σ, and sensitivity gain coefficients obtained for NLISA vs. standard ELISA for insulin, IFN-γ, and p24. *LOD_2_σ.

## Conclusions

Current ultrasensitive detection techniques in the fM range or below are complex and the increase in sensitivity comes at the expense of increased cost and with the requirement of expensive infrastructure and qualified personnel. We here proposed a novel approach, NLISA, which combines simplicity, ease of use, and a broad dynamic range (4-5 orders of magnitude) of quantitative detection while attaining (sub)-fM sensitivity. We exploit the unique properties of 38-nm YVO_4_:Eu nanoparticles obtained through a metavanadate synthesis route and subsequently annealed at 200°C to improve crystallinity. Their ultrahigh extinction coefficient and their 18% quantum yield lead to extremely bright probes. A simple home-made reader ensures excitation with a 275-nm LED and readout in a 96-well plate format using a photomultiplier.

This technology is easily generalizable: it can be applied to any polypeptide or protein provided the corresponding biorecognition molecules are available. Biorecognition molecules can either be directly coupled covalently to the NH_2_- or COOH-functionalized nanoparticles or can be biotinylated and coupled to streptavidin-coated nanoparticles for increased versatility.

This approach combining ultrasensitive (sub)-fM, quantitative detection with straightforward detection should push forward earlier diagnosis and treatment for numerous diseases, enable less invasive diagnostics in human samples containing biomarkers in low concentrations, and favour the emergence of novel biomarkers. The simplicity of this method based on a transportable reader should facilitate its implementation in low-infrastructure areas of the world. Moreover, NLISA performances may be further enhanced using label engineering for the controlled formation of nanoparticle complexes.

## Acknowledgments

We are grateful to Pascal Preira and Max Richly for their participation in the early stages of this work and to Xavier Solinas for his expert help with the reader electronics. We acknowledge funding from the Centre Interdisciplinaire d’Etudes pour la Défense et la Sécurité (CIEDS), projects DAVID and DOMUS.

## Materials and methods

### Nanoparticle synthesis, annealing, and functionalization

#### Synthesis of YVO_4_:Eu^3+^ nanoparticles

The particles synthesis was adapted from the V3 protocol of our previous work^29^. A solution was prepared with 0.08 M Y(III)(NO_3_)_3_ (Sigma, 99.9%) and 0.02 M Eu(III)(NO_3_)_3_ (Sigma, 99.8%), in 25 mL Milli-Q water. This solution was quickly added to a solution of 0.1 M NaVO_3_ (Sigma, 98%), and 0.3 M NaOH in 25 mL Milli-Q water under vigorous stirring. The obtained mixture was left under stirring for 14 days and then purified 3 times by centrifugation (26323 g for 25 min) followed each time by redispersion in 50 mL of Milli-Q water using an ultrasonic probe sonicator (Fisher, model: 7501D, 130W).

Surface silicatation of the particles was achieved by adding sodium silicate (Merck, [Si]= 6.07M, Na_2_O: 7.5-8.5%, SiO_2_: 25.5-28.5%) with an amount leading to a Si/V molar ratio of 10. The solution was then left under stirring for 3 hours and further purified 3 times by centrifugation (26323 g, 25 min) followed each time by redispersion in 50 mL of Milli-Q water using an ultrasonic probe sonicator. The colloidal suspension of silicated nanoparticles was then annealed at 200°C in a µ-wave oven (CEM Discover SP + Explorer) for 2 hours.

After that treatment, a second step of silicatation was achieved following the same protocole as the first time: sodium silicate was added with an amount leading to a Si/V molar ratio of 10. The solution was then left under stirring for 3 hours and then purified 3 times by centrifugation (26323 g, 25 min) followed by dispersion in 50 mL of Milli-Q water using an ultrasonic probe sonicator.

The obtained suspension of YVO_4_:Eu^3+^ nanoparticles (50 mM in vanadate ions) was homogeneous and slightly light scattering. For the purpose of our experiments, this solution was diluted 20 times in a pH7 buffer (VWR). In this final colloidal suspension, the particle size measured by DLS (Malvern Zetasizer) was found to be 73 nm (in number, PDI 0.1) and the particle zeta potential was found to be -49 mV.

### Functionalization of the nanoparticles

Nanoparticles were functionalized according to previous work^34^ to enable their covalent coupling with streptavidin. To this end, 75 mL of re-silicated YVO_4_:Eu^3+^ nanoparticles at 3 mM in vanadate ions were added dropwise (1 mL min^-1^) to 225 mL of boiling ethanol containing 265 µL of APTES. The solution was left under reflux for 24 hours at 92 °C to ensure a uniform cross-linked aminated network surrounding the particles. The solids were then separated by centrifugation (26323 g for 30 min), washed two times using an ethanol/water mixture (3/1 v/v), and redispersed in 5 mL of Milli-Q water. Finally, the colloidal suspension was added dropwise to 20 mL of succinic anhydride in DMF (0.25 g mL^-1^) and stirred overnight at room temperature. The particles were purified by 3 cycles of centrifugation/redispersion in the ethanol/water mixture and resuspended in 5 mL of MES buffer (50 mM, pH 5 to 6).

### NP bioconjugation step

The appropriate volume was pipetted to obtain 150 µL of –COOH coated nanoparticles at a concentration of 200 nM. The sample was then centrifuged 15 min at 13700 g (Servall Legend Micro17R Centrifuge) and the supernatant is removed. 1-ethyl-3-(3-dimethylaminopropyl) carbodiimide (EDC) and N-hydroxy succinimide (NHS) were solubilized at a concentration of 50 mg.mL^-1^ each in a 2-(N-morpholino)ethane sulfonic acid (MES) 50 mM buffer at pH 5.5. This solution was quickly added to the nanoparticle pellet and the sample was sonicated 15 s at 50% power (Cole-Parmer Model CV18). The sample was left under stirring at room temperature during 25 min and was then centrifuged 15 min at 15000 g to remove EDC/NHS. The nanoparticles were then redispersed in 50-mM phosphate buffer pH 7.4 by sonication during 15 s at 50% power. 40 equivalents of streptavidin (Sigma, s4762-10MG), i. e. 40 streptavidin molecules per nanoparticle were then added and the sample was incubated at 25 °C under stirring at 800 rpm (Eppendorf Thermomixer C) for 2h30. The sample was centrifuged 15 min at 13700 g to remove unbound streptavidin molecules and the pellet was resuspended in 500 µL blocking buffer (50-mM phosphate buffer pH 7.4 containing 2% mPEG-NH_2_ 500 (Biochempeg, ref. MF001005-500) by pulsed sonication for 10 s by alternating 1s on/1s off at 20% power in ice. The sample was then left under stirring for 1 h at room temperature and centrifuged 15 min at 13700 g to remove unbound PEG molecules. The supernatant was removed and the streptavidin-nanoparticle conjugates were redispersed in conservation buffer (Tris 20 mM, pH 8, 1% BSA) and stored at -80°C. The final nanoparticle concentration obtained was close to the initial one.

### Dosage of the conjugated streptavidin-NP ratio

We used a Bradford assay (BioRad 5000006) performed on the supernatant after the centrifugation step to remove unbound streptavidin molecules to deduce the number of streptavidin molecules actually bound per nanoparticle. We tested different SA:NP ratios and found that the optimal sensitivity was obtained for a 40:1 ratio in the coupling protocol (Supp. Fig. S1). A coupling efficiency between 80 and 90% was obtained, i. e. typically between 37:1 and 32:1.

### NLISA reader

The excitation module was composed of a 100-mW, 275-nm LED (Hex-S6060-DR250-W275-P100-V6.5, Laser Components), pulse-width modulation controlled with a voltage input module NI-9215, cooled down by a 10-mm heatsink associated to a 5-V fan. The UV beam went through a ∼1 mm-hole to avoid the excitation of neighboring wells permitting a single well excitation without crosstalk (Supp. Table S1). The detection module included a collecting lens with a 0.79 numerical aperture (ACL25416U-A, Thorlabs), a f=100 mm focusing lens (LA1509-A, Thorlabs), a 45° mirror (MRA25-E02, Thorlabs), and two 620-nm filters (FF01-620/14-25, Semrock) inserted in front of a 300-800 nm photomultiplier module (PMM02, Thorlabs).

The readout of the multiwell plate was done with 2 stepper motors (17HS15-1504S-X1), a lead screw in the Y direction and a pulley-belt system in the X direction. The XY translations were managed by an Arduino UNO board with the grbl 1.1 library upload (Github) complemented by a CNC shield. Most of the mechanical assembly parts were 3D-designed and printed with polylactic acid or CPE+ (Ultimaker) black filaments.

Concerning the signal acquisition, each microplate well was excited by the UV LED with a total power of 2 mW measured at the well bottom, during 900 ms, and we recorded the voltage readout of the photomultiplier module with a voltage output module (NI-9269) with a frequency of 100 kHz. We averaged the last 600 ms of acquisition (after 300 ms to allow for the LED pre-heating). We did not observe any saturation effects of the PMT in all the experiments performed. The communications with all electronic parts and data analysis were managed by a homemade Labview program.

The detection efficiency of this reader is 𝜂_𝑟𝑒𝑎𝑑𝑒𝑟_ = 0.59%. With respect to the microscope setup, this 35x weaker detection efficiency is mainly due to the lower quantum yield of the photomultiplier tube (at 617 nm, 0.04 vs. 0.93 for the EM-CCD camera) and to the smaller solid angle of the emission collection system (the fraction of the total emission volume detected is 0.19 vs. 0.31 for the microscope objective).

### Patient sample preparation and characterization

The study was conducted in accordance with the Declaration of Helsinki. All patients gave written informed non-opposition for a de-identified electronic version of their medical records to be used for research purposes and for access to the remains of blood following standard follow-up for research purpose. The study was approved by the Scientific Committee of Ile de France Paris VII.

Samples of blood from people leaving with HIV (PLWH) were anonymized in the clinical services and transferred to the immunomonitoring laboratory. Plasma is collected following whole blood centrifugation at room temperature during 10 minutes at 244 g (Rotanta 460R, Hettich), frozen and stored at -80°C. Peripheral blood was also collected for evaluation of immune parameters. Plasma HIV-l RNA viral load is measured by a routine assay (with a detection limit of 200 copies/mL) and ultrasensitive (US) assays as described in Ref.^66^. Ultrasensitive HIV-RNA quantifications (US HIV-RNA) were performed using the Generic HIV real-time PCR assay (Biocentric, Bandol, France). The threshold for US assays ranged from 1 to 40 copies/mL, depending on the available plasma volume (0.5 to 15 mL). For subsequent analyses performed in BSL1 laboratory, a step of inactivation of plasma was performed during 15 minutes using formaldehyde solution (Sigma, Merck, USA) to reach a final concentration of 1% of formaldehyde solution.

