## supporting information for "Ultrasensitive quantitative protein detection using Eu-ion doped vanadate nanoparticles"

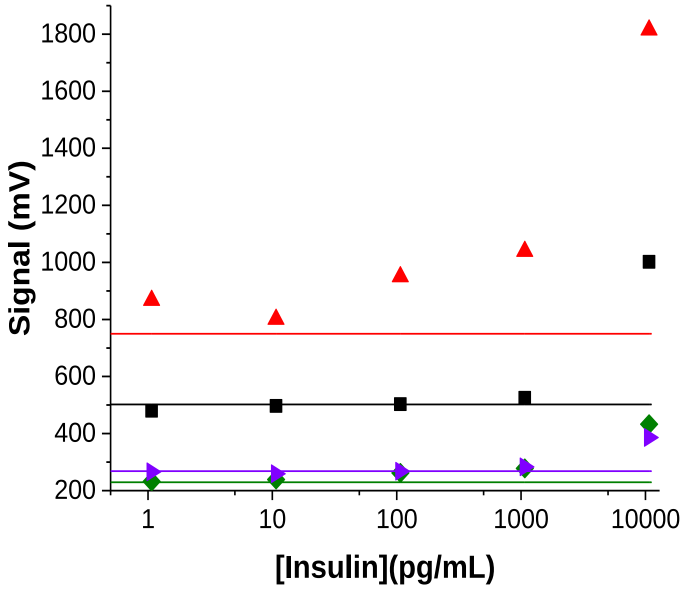


**Fig. S1.** Optimization of the number of streptavidin molecules per nanoparticle. Insulin detection was performed for 36:1 (red), 20:1 (black), 53:1 (green) and 80:1 (purple) streptavidin-nanoparticle conjugates. Full lines indicate the $LOD_{3\sigma}$ for each condition. The figure shows that the optimal sensitivity is obtained for the 36:1 streptavidin-nanoparticle conjugates.

| | | | | | |
|---|---|---|---|---|---|
| 54.35 | 2216.38 | 61.93 | 2727.31 | 63.86 | 55.72 |
| 2509.84 | 60.71 | 2475.75 | 65.12 | 2257.66 | 56.19 |
| 55.38 | 1780.11 | 56.35 | 63.68 | 65.83 | 68.08 |
| 55.85 | 54.89 | 55.48 | 55.15 | 55.36 | 55.61 |
| 55 | 53.98 | 55.98 | 54.98 | 59.39 | 55.61 |
| 56.31 | 55.8 | 55.99 | 65.37 | 55.54 | 55.46 |
| 59.47 | 56.32 | 56.41 | 56.13 | 56.12 | 55.85 |
| 55.85 | 54.37 | 56.07 | 55.61 | 55.61 | 55.43 |

| | | |
|---|---|---|
| Mean adjacent wells | 59.6 | ±4.8 |
| Mean far away wells | 56.4 | ±2.3 |

**Table S1.** Cross talk verification. Top: Signals in mV obtained for 48 wells of a 96-well plate. In red/pink, wells that have been incubated with a high concentration of nanoparticles yielding a signal similar to the highest signals observed in the experiments; in yellow, wells without nanoparticles adjacent (or diagonally in contact) to wells with a high signal due to nanoparticles; in green, wells without nanoparticles lying further away from wells yielding a strong nanoparticle signal. Bottom: The mean signal value and standard deviation from wells without nanoparticles adjacent or in contact with wells yielding a strong nanoparticle signal (yellow) and from wells without nanoparticles lying further away from wells with nanoparticles. The overlap between these two types of signal values shows a negligible crosstalk between adjacent wells. In addition, this figure confirms the absence of variability between wells located at different positions on the multiwell plate.

#### Linearity of the reader photomultiplier signal

We used a strongly attenuated HeNe laser at different power levels to investigate the linearity between the incident laser power and signal in mV measured by the photomultiplier tube (PMT). A linear relationship was found for the whole range of measured signals.

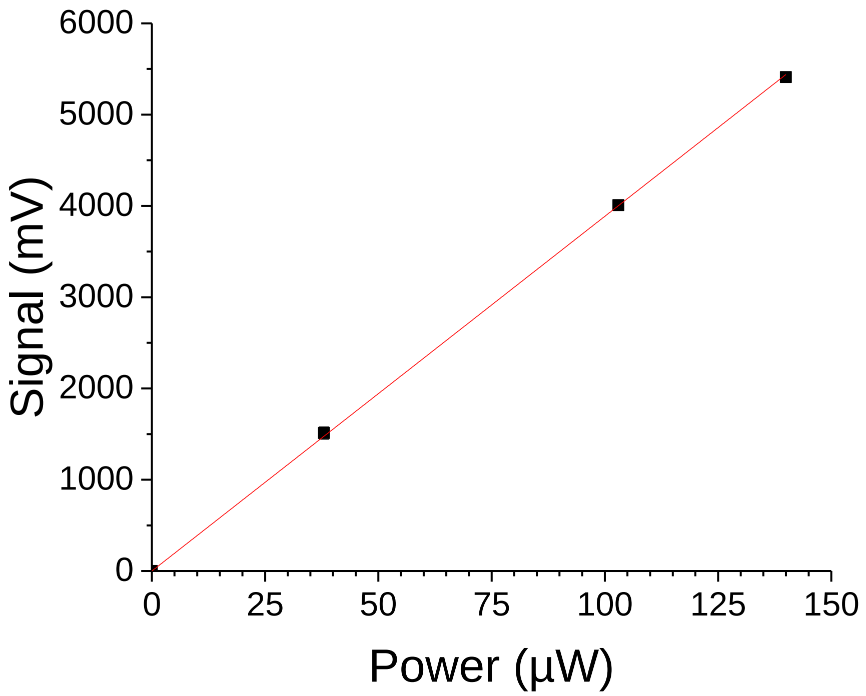


**Fig. S2.** Linear relationship between the incident optical power at 632.8 nm and the signal measured by the PMT.

| Insulin Concentration (pM) | Max Insulin attached | Mean PM signal-blank (mV)* | Nanoparticles/well | Ratio Nanoparticles/ Max Insulin attached |
|---|---|---|---|---|
| 8.34 | $5.0\times10^{8}$ | 1201 ± 65 | $2.0\times10^{6} \pm 1.07\times10^{5}$ | $4\times10^{-3} \pm 2.1\times10^{-4}$ |
| 0.834 | $5.0\times10^{7}$ | 1003 ± 35 | $1.7\times10^{6} \pm 5.76\times10^{4}$ | $0.03 \pm 1.1\times10^{-3}$ |
| $8.34\times10^{-2}$ | $5.0\times10^{6}$ | 764 ± 58 | $1.3\times10^{6} \pm 9.57\times10^{4}$ | $0.25 \pm 1.9\times10^{-2}$ |
| $8.34\times10^{-3}$ | $5.0\times10^{5}$ | 516 ± 31 | $8.5\times10^{5} \pm 5.14\times10^{4}$ | 1.70 ± 0.10 |
| $8.34\times10^{-4}$ | $5.0\times10^{4}$ | 204 ± 22 | $3.4\times10^{5} \pm 3.66\times10^{4}$ | 6.7 ± 0.73 |

**Table S2.** Calculation of the number of nanoparticles labelling each captured insulin molecule based on the PMT signal of the reader and considering that all the insulin molecules in solution have been captured. *experiments done in triplicate.

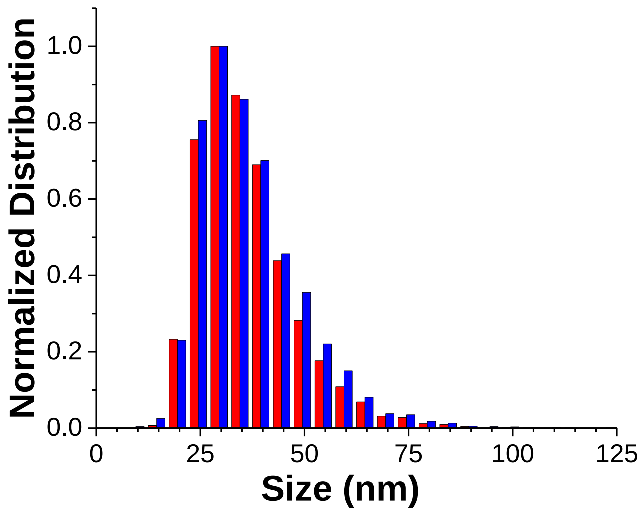


**Fig. S3.** Comparison between the size distribution of annealed $YVO_4$:Eu 20% nanoparticles (in red, same figure as in Fig. 1D) and the size distribution of NP-SA conjugates (in blue, same histogram as the histogram in blue in Fig. 4B). The size distributions were obtained from the number of detected photons using wide-field microscopy (see main text). The overlap of the two histograms shows that the functionalization does not modify the nanoparticle size.

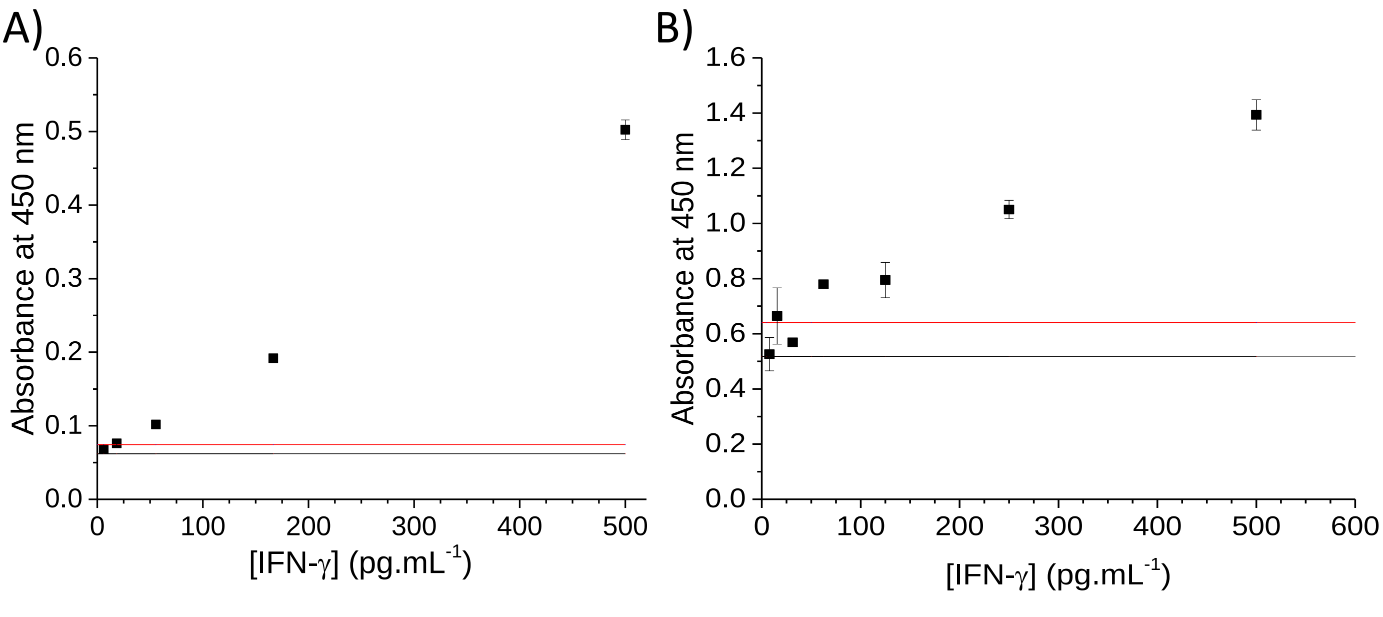


**Fig. S4.** IFN-γ detection using the commercial ELISA kit which measures the absorbance at 450 nm A) in buffer and B) in undiluted serum. Experiments were performed in triplicate. The error bars correspond to the standard error on the mean. The black (red) lines indicate the signal corresponding to the mean value (mean value plus three standard deviations) of 10 blank samples. We took as $LOD_{3\sigma}$ the concentration corresponding to the crossing point between the red line and the line connecting the last data point below the red line as a function of increasing concentration and the consecutive data point above the red line.

| IFN-γ Concentration (pM) | Max IFN-γ attached | Mean PM signal-blank (mV)* | Nanoparticles/well | Ratio Nanoparticles/ Max IFN-γ attached |
|---|---|---|---|---|
| 29 | $1.8x10^{9}$ | 248 ± 15 | $4.1x10^{5} \pm 2.4x10^{4}$ | $2.3x10^{-4} \pm 1.4x10^{-5}$ |
| 5.8 | $3.5x10^{8}$ | 184 ± 8.4 | $3.0x10^{5} \pm 1.4x10^{4}$ | $8.7x10^{-4} \pm 4.0x10^{-5}$ |
| 1.2 | $7.0x10^{7}$ | 85.6 ± 8.6 | $1.4x10^{5} \pm 1.4x10^{4}$ | $1.9x10^{-3} \pm 2.0x10^{-4}$ |
| $2.3x10^{-1}$ | $1.4x10^{7}$ | 48.3 ± 4.4 | $8.0x10^{4} \pm 7.3x10^{3}$ | $5.7x10^{-3} \pm 5.2x10^{-4}$ |
| $4.7x10^{-2}$ | $2.8x10^{6}$ | 38.6 ± 4.2 | $6.4x10^{4} \pm 6.9x10^{3}$ | $2.3x10^{-2} \pm 2.5x10^{-3}$ |
| $9.3x10^{-3}$ | $5.6x10^{5}$ | 27.2 ± 1.9 | $4.5x10^{4} \pm 3.2x10^{3}$ | $8.0x10^{-2} \pm 6.0x10^{-3}$ |

| IFN-γ Concentration (pM) | Max IFN-γ attached | Mean PM signal-blank (mV)* | Nanoparticles/well | Ratio Nanoparticles/ Max IFN-γ attached |
|---|---|---|---|---|
| 29 | $1.8x10^{9}$ | 72.9 ± 3.6 | $1.1x10^{5} \pm 5.9x10^{3}$ | $6.3x10^{-5} \pm 3.4x10^{-4}$ |
| 5.8 | $3.5x10^{8}$ | 58.1 ± 11 | $8.7x10^{4} \pm 1.9x10^{4}$ | $2.5x10^{-4} \pm 5.4x10^{-5}$ |
| 1.2 | $7.0x10^{7}$ | 38.0 ± 4.2 | $5.4x10^{4} \pm 6.9x10^{3}$ | $7.7x10^{-4} \pm 1.0x10^{-4}$ |
| $2.3x10^{-1}$ | $1.4x10^{7}$ | 24.5 ± 3.4 | $3.1x10^{4} \pm 5.6x10^{3}$ | $2.2x10^{-3} \pm 4.0x10^{-4}$ |
| $4.7x10^{-2}$ | $2.8x10^{6}$ | 16.1 ± 3.0 | $1.8x10^{4} \pm 5.0x10^{3}$ | $6.2x10^{-3} \pm 1.8x10^{-3}$ |

**Table S3.** Calculation of the number of nanoparticles labelling each captured molecule IFN-γ based on the PMT signal of the reader and considering that all the IFN-γ molecules in solution have been captured. Data in buffer (top) and in undiluted serum (bottom). *experiments done in triplicate

| p24 Concentration (pM) | Max p24 attached | Mean PM signal-blank (mV)* | Nanoparticles/well | Ratio Nanoparticles/ Max p24 attached |
|---|---|---|---|---|
| 16,8 | $1.01x10^{9}$ | 88.4 ± 5.6 | $1.5x10^{5} \pm 9.2x10^{3}$ | $1.4x10^{-4} \pm 9.1x10^{-6}$ |
| 1,68 | $1.01x10^{8}$ | 57.3 ± 7.0 | $9.5x10^{4} \pm 1.2x10^{4}$ | $9.4x10^{-4} \pm 1.1x10^{-4}$ |
| $1.68x10^{-1}$ | $1.01x10^{7}$ | 27.2 ± 3.1 | $4.5x10^{4} \pm 5.1x10^{3}$ | $4.4x10^{-3} \pm 5.1x10^{-4}$ |
| $1.68\ x10^{-2}$ | $1.01x10^{6}$ | 18.2 ± 3.5 | $3.0x10^{4} \pm 5.8x10^{3}$ | $3.0x10^{-2} \pm 5.7x10^{-3}$ |

**Table S4.** Calculation of the number of nanoparticles labelling each captured p24 molecule based on the PMT signal of the reader and considering that all the p24 molecules in solution have been captured. *experiments done in triplicate.

**Supporting Materials and Methods**

**NP characterization**

Transmission Electron Microscopy (TEM) analysis was carried out using a JEOL JEM-2010F microscope with the following specifications: high tension (HT) objective lens Cs = 1.4 mm, Cc = 1.8 mm, Focal length = 2.7 mm. The electron microscope was operated at 200 kV and was equipped with a Gatan US4000 CCD camera (Fig. 1A-B). 20 µL of diluted particle suspensions were deposited on a Parafilm sheet. Carbon-coated copper grids (200 mesh), previously treated with a positive glow-discharge were brought into contact with the sample droplets during 1 minute. Subsequently, the grids were removed and washed sequentially with two droplets (20 µL each) of Milli-Q water, with the excess fluid being removed by placing a filter paper on the grid's edge. The grids were then air-dried under ambient conditions. Image analysis was performed using ImageJ software with manual selection. The average nanoparticle full width and length were found to be 31.1 and 43.8 nm, respectively. Assuming that the third axis is identical to the short axis, we could calculate the volume of the nanoparticles and then deduce the diameter of a spherical particle with equal volume (see size distribution in Fig. 1D).

Additionally, Dynamic Light Scattering (DLS) and ζ potential profiles were obtained using a Malvern Zetasizer Nano ZS. The measurements were conducted on nanoparticle suspensions (0.5 mM in vanadate ions) in Milli-Q water after homogenization with an ultrasonic processor

(80 W, 20 Hz, 2 min). The ζ potential was measured after each functionalization step and changed as expected (Fig. S4).

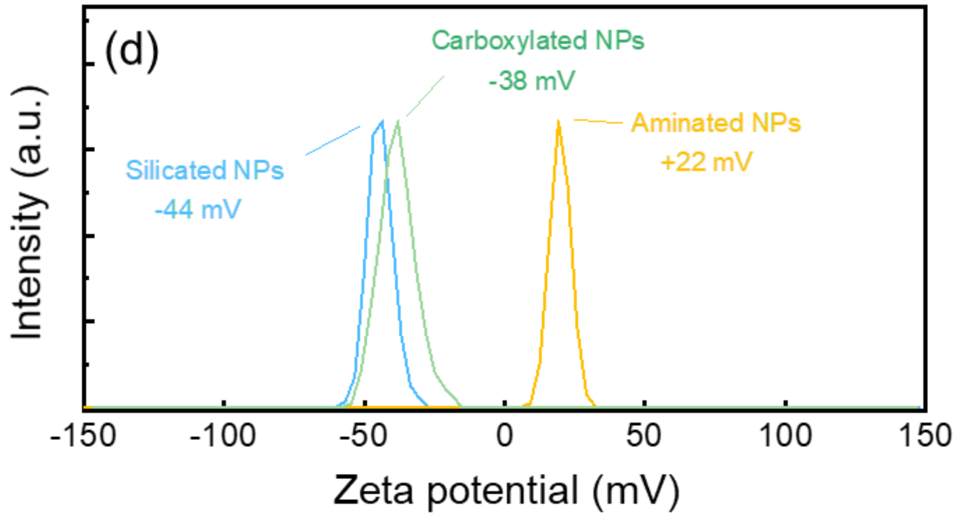


**Fig. S5.** Zeta potential of the nanoparticles after the different functionalization steps: after silicatation (blue), amination (yellow), and carboxylation (green).

The luminescence overall quantum yields q of particle suspensions were estimated using a Horiba Quanta φ integrating sphere and rhodamine 6G solution ($10^{-5}$ M in ethanol, $q_{Rho}$ = 0.94)[64] as an internal standard having the same optical density (OD ~ 0.2) and excited at the same wavelength (280 nm).

### Determination of nanoparticle concentration

Nanoparticles were dissolved in acid and the vanadate ion concentration was measured by a colorimetric approach as follows[24]. 90 µL of nanoparticle solution were diluted in 1 mL $H_2O$ milliQ in triplicate. The sample was then centrifuged 15 min at 13700 g (Servall Legend Micro17R Centrifuge). The pellet was dissolved in 100 µL of HCl 12 N and left 20 min at room temperature (RT). 400 µL of $H_2O$ were added and the solution was transferred to a glass dish containing 3 mL of HCl 1.15 N. The sample was then incubated 45 min at 75 °C and was subsequently left to cool down 15 min at RT. Then, 52.5 µL of $H_2O_2$ were added and the solution was left 5 min at RT. The absorbance at 800 nm and 405 nm was then read. The vanadate concentration $C$ was calculated by: $C\ (mM) = \frac{(A_{405} - A_{800}) * D}{\varepsilon_{405}}$, where $D$ is the vial thickness, and $\varepsilon_{405}$ the extinction coefficient at 405 nm of the vanadium complexes formed.

The nanoparticle concentration could be deduced from the concentration in vanadate ions based on the mean nanoparticle volume and on the number of vanadate ions per nanoparticle as was explained in Ref[19]. The number of vanadate ions for a given nanoparticle volume is calculated from the unit cell volume and from the fact that there are 4 vanadate ions per unit cell (see main text).

**Optical microscopy**

Several suspensions of streptavidin-conjugated $YVO_4:Eu^{3+}$ nanoparticles (830 pM to 830 fM) (56 µM to 56 nM in vanadate ions) were prepared in PBS buffer at pH 7.4 and 100 µL of each were deposited in multiwell plates with a 190-µm thick cyclo-olefin well bottom (Greiner Screenstar, Dutscher 655866) in duplicate. The plate was incubated for 48 hours at 4°C under stirring at 300 rpm (Eppendorf Thermomixer C) and washed five times with PBS buffer at pH 7.4. The signal due to non-specific binding of the nanoparticles at the well bottom was read on the homemade UV microplate reader. Then each well was observed under a wide-field microscope for single-particle observation (Olympus IX83 equipped with a plan-apochromat, 63x, 1.4 N. A. Zeiss objective Ref. 440762 and a Photometrics Evolve EM-CCD 512x512 camera) using excitation with a 466-nm diode laser (Modulight). We used 466-nm excitation because current high numerical-aperture microscope objectives for wide-field single-molecule imaging do not transmit sufficiently in the UV to allow excitation of the nanoparticles via the strong vanadate matrix absorption. We recorded images and analysed five different regions per well with a homemade MATLAB program to count the number of single nanoparticles visible in each image and the photon number per single particle emission spot.

We consider the nanoparticle quantum yield upon excitation of the vanadate matrix to be the same as that upon direct $Eu^{3+}$-ion excitation under the optical microscope because the non-radiative losses are mainly due to the non-radiative processes taking place during the last $^5D_0$-$^7F_2$ deexcitation step.

**NLISA insulin detection and quantification**

We used the Abcam insulin kit (Ref. ab100578) as indicated by the supplier by replacing the horseradish-peroxidase labeled detection antibodies with biotinylated detection antibodies that we used for binding to the streptavidin-nanoparticle conjugates. The kit contains recombinant insulin, diluent solutions and a microwell plate coated with capture antibodies, biotinylated detection antibodies, and the chimer streptavidin-HRP.

An insulin solution is prepared in diluant A to obtain a 10 $ng.mL^{-1}$ concentration. Then, the solution is diluted in cascade by a factor of 10 (triplicate for each concentration and fifteen blanc solutions) and 100 µL solutions of various insulin concentrations are added to the microplate wells. The microplate is then incubated 2h30 at 25°C under stirring at 300 rpm (Eppendorf Thermomixer C).

In parallel, streptavidin-nanoparticle conjugates are incubated with 60 equivalents of biotinylated detection antibody in phosphate-buffered-saline (PBS) pH 7.4 for 1h30 at 25°C. The solution is then centrifuged 15 minutes at 13 000 g to remove unbound antibodies. The pellet is resuspended in diluant B by pulsed sonication 10 s 1s on/1s off on ice to obtain a final 4.5-nM nanoparticle concentration.

The commercial microwell plate is then washed three times with the washing buffer supplied with the kit. 100 µL of nanoparticle-antibody conjugate solution is added per well and the plate is incubated 1h30 at 25°C under stirring at 300 rpm. The plate is washed three times with the washing buffer and one time with PBS pH 6.6. Finally, 200 µL of PBS pH 6.6 are added to each well and the plate is read with the homemade reader.

**NLISA IFN-γ detection and quantification**

We used the commercial kit Thermofisher Ref. 88-7316-88 containing recombinant IFN-γ, diluents, capture antibodies, biotinylated detection antibodies, and the chimer streptavidin-HRP.

The provided capture antibody solutions were diluted 250 times in the coating buffer of the kit and 100 µL was incubated in each well of a multiwell plate (Greiner 655097) for 16 h at 4°C under stirring at 300 rpm (Eppendorf Thermomixer C). The wells were then washed two times with Elisa/Elispot diluent provided with the kit. Then 200 µL/well of Elisa/Elispot diluent was added during 2 h at 4°C under stirring at 300 rpm as a blocking step.

A range of IFN-γ concentrations was prepared (250 $pg.mL^{-1}$ to 25 $fg.mL^{-1}$) and 100 µL of the various IFN-γ solutions were added per well. The plate was then incubated 16 h at 4°C, 300 rpm. The wells are then washed three times with PBS pH 7.4.

In parallel, streptavidin-nanoparticle conjugates were mixed with 60 equivalents of detection antibody (4S.B3 Thermofischer 13-7319-85) in PBS pH 7.4 during 1h at room temperature under gentle stirring. The solution was then centrifuged 15 minutes at 13 700 g. The pellet was suspended in PBS pH 7.4 by pulsed sonication 10 s (1s on/1s off) on ice to obtain a final 4.5 nM nanoparticle concentration.

100 µL of the nanoparticle-detection antibody solution was added in each well and the plate was incubated 1h30 at 25°C under stirring at 300 rpm. The plate was then washed five times with 200 µL PBS pH 7.4, 200 µL PBS pH 7.4 was added, and the plate was read with the homemade reader.

**NLISA HIV GAG-p24 detection and quantification**

We used commercial recombinant p24 capture and detection antibodies (Bio-techne, NBP3-06466, and Mab73602, respectively).

**Antibody biotinylation.** We used the biotinylation kit Bio-techne 370-0010 to biotinylate the detection antibodies. We added 10 µL of Modifier reagent to 100 µL of capture antibody solution at 1 mg.mL-1 in PBS pH 7.2. We then added this solution directly to 50 µg of lyophilized biotin mix leading to 3.5 of biotin equivalents in the final solution. We resuspended gently and the solution was wrapped in aluminium foil at room temperature during 30 minutes. We then added 10 µL of Quencher reagent for every 10 µL of antibody used and mixed gently.

**NLISA.** We diluted the capture antibody to a concentration of 10 µg.mL-1 in PBS and immediately coated a 96-well microplate ( Greiner 655097) with 100 µL per well. Following the supplier protocol, we sealed the plate and incubated overnight at room temperature. We then washed three times by removing the solution from each well and washing with wash buffer PBS pH 7.2, Tween 0.05%. We finally filled each well with wash Buffer (400 µL). We then blocked the wells by adding 300 µL PBS pH 7.2, BSA 2% to each well and incubating at room temperature for a minimum of 1 hour. The washing step above (three times) was then repeated.

p24 solutions were prepared at concentrations between 150 and 300 ng/mL. 100 µL of the different concentration solutions were added per well in triplicate and 100 µL of PBS buffer was added for the blank measurements. We then incubated 2 hours at room temperature. The wells were washed three times with PBS pH 7.2, Tween 0.05%.

In parallel, nanoparticle-streptavidin conjugates were mixed with 60 equivalents of the biotinylated detection antibody in PBS pH 7.2 during 1h at room temperature under gentle stirring. The solution was then centrifuged 15 minutes at 13 700 g. The pellet is suspended in

PBS pH 7.2 by pulsed sonication 10 s (1s on/1s off) on ice to obtain a final 4.5 nM nanoparticle concentration.

100 µL of the resulting nanoparticle-detection antibody solution are added to each well, and the plate is incubated 1h30 at 25°C under stirring at 300 rpm. The plate is then washed four times with PBS pH 7.2, Tween 0.05% and once with PBS pH 7.2. 200 µL PBS pH 7.2 is added to each well and the plate is read with the homemade reader.

**ELISA measurements**

The ELISA measurements for insulin and IFN-γ were performed following the supplier protocol (see kit refs. above).
